# The Fossilized Birth Death Process with heterogeneous diversification rates unravels the link between diversification and specialisation to a carnivorous diet in Nimravidae (Carnivoraformes)

**DOI:** 10.1101/2025.07.15.664897

**Authors:** Nils Chabrol, Hélène Morlon, Joëlle Barido-Sottani

## Abstract

Bayesian phylogenetic inference uses more and more complex diversification models as tree priors to test new macroevolutionary hypotheses. However, those models are usually developed in a neontological framework, despite the increasing number of datasets covering both extant and fossil taxa, as well as the fact that many clades are entirely extinct. In this paper, we develop the F-ClaDS model, a Fossilized-Birth-Death (FBD) version of the cladogenetic diversification rate shift (ClaDS) model, in BEAST2. ClaDS estimates partially inherited branch-specific rates from a phylogeny, providing a nuanced and detailed perspective of the variations in diversification across the tree. Our extension allows the integration of fossil samples directly into the phylogeny. We apply our new implementation to a dataset of 36 Nimravidae, a fully extinct carnivoraform clade that spanned from the Early Eocene to the Late Miocene, which species had different degrees of specialisation to a carnivorous diet. We show that using different tree priors does not affect substantially the topology of the inferred trees, but affects the ages of nodes and tips, as well as branch-lengths. F-ClaDS also recovers more species as sampled ancestors than the homogeneous FBD model. The branches with the highest speciation and extinction rates are those corresponding to the hypercarnivorous clades (*Hoplophoneus* and the barbourofelins), supporting the view that specialization to a hypercarnivorous diet can spur speciation, but also increase extinction risk, especially during times of global ecosystem change, potentially due to a high position in the trophic chain.

## 1 Introduction

Estimating the speciation and extinction rates of species is a major goal of macro-evolution studies. These rates can be used to explain global geographical diversity patterns like the Latitudinal Diversity Gradient (LDG) (Mittelbach et al., 2007; Rolland et al., 2014), to assess the deep-time history of a clade (Alfaro et al., 2009; Jetz and Pyron, 2018) or to correlate diversification dynamics with environmental parameters (Velasco et al., 2016; Kong et al., 2017) or trait evolution (Rabosky et al., 2013; Cantalapiedra et al., 2014). A widely used set of methods that allows estimating diversification rates rely on birth-death models. These methods estimate rates of “birth” (i.e. speciation, usually noted *λ*) and “death” (i.e. extinction, usually noted *µ*) from time-calibrated phylogenetic trees. While initial developments in the field assumed homogeneous rates (i.e., that both *λ* and *µ* are identical across branches in the tree), studies based on the shape of reconstructed trees have unambiguously shown that trees are generally more asymmetric than what is expected under homogeneous diversification rates (Guyer and Slowinski, 1991; Aldous, 1996). To address this issue, several birth-death processes with heterogeneous diversification rates have been developed (Alfaro et al., 2009; Morlon et al., 2011; Rabosky et al., 2013; Barido-Sottani et al., 2020; Quintero et al., 2024). In particular, Maliet et al. (2019) developed the cladogenetic diversification rate shift model, ClaDS, a birth-death process that allows small changes in *λ* and *µ* at each speciation event. The specificity of ClaDS is to explicitly model progressive rate variations on all branches, while previous models, often based on the multi-type birth-death process, assumed punctual regime shifts. Punctual regime shifts capture major ecological transitions in the tree, or the appearance of a key trait leading to an adaptive radiation (Stroud and Losos, 2016), but not the complex interplay between species evolving ecologies and their specific spatial and environmental context (Morlon et al., 2024); models with many small shifts in diversification rates, seem to be better supported on empirical data (Ronquist et al., 2021). Maliet et al. (2019) initially fitted the ClaDS model in a Monte Carlo Markov Chain (MCMC) Bayesian framework where the computation of the likelihood required solving Ordinary Differential Equations (ODEs). Later, Maliet and Morlon (2022) developed an algorithm based on Bayesian Data Augmentation (DA) that makes the inference more efficient computationally and provides augmented trees, which complement the reconstructed tree with probable lineages that are unobserved because of extinction or incomplete sampling.

Beside their use on fixed trees, birth-death models of diversification have been widely used in Bayesian phylogenetic inferences as a tree prior, informing the tree topology and branching times, and allowing to jointly estimate the tree and the diversification rates of the studied clade from sequence alignments (Rannala and Yang, 1996; Höhna et al., 2016; Bouckaert et al., 2019). In the family of birth-death models with heterogeneous diversification rates, this so-called “full phylogenetic inference” approach has been implemented for the multitype birth-death process (Barido-Sottani et al., 2020), as well as for ClaDS in its DA version (Barido-Sottani and Morlon, 2023). Both models are implemented in BEAST2.

Another area of development in birth-death models concerns the integration of fossil data. Under “node-dating” approaches, fossils are used to calibrate internal nodes in phylogenetic Bayesian inference methods (Near and Sanderson, 2004), but are not accounted for when inferring the diversification process. This changed with the development of the Fossilized Birth-Death process (FBD) (Stadler, 2010): this model directly incorporates fossils in the phylogeny by introducing an additional parameter *ψ*, which captures the rate of both fossilization and recovery of fossils by paleontologists. Fossils can occur as distinct lineages, referred to as “fossil tips” or on already existing branches, as ancestors of already represented extinct or extant taxa, referred to as “sampled ancestors”. The FBD process is now increasingly used as a prior for estimating divergence times in Bayesian phylogenetic “total evidence” analyses, which use morphological characters to place fossils (Zhang et al., 2016; Gavryushkina et al., 2017; Matschiner et al., 2017). Initially implemented with homogeneous diversification and fossilization rates (Heath et al., 2014), the FBD model has now been implemented for the multi-type birth-death process while accounting for heterogeneous fossilization rates (Barido-Sottani and Morlon, 2025), as well as for the birth-death-diffusion model, although with a fixed tree (Quintero et al., 2024). In the context of full phylogenetic inference, Barido-Sottani and Morlon (2025) showed that incorporating fossils with informative characters significantly improves the accuracy of both the phylogeny and diversification rates.

Here, we implemented a FBD version of ClaDS (F-ClaDS) for full phylogenetic inference, building upon its DA implementation in the ClaDS package in BEAST 2 (Barido-Sottani and Morlon, 2023); the implementation is available in that package. Previous work has shown that diversification rate estimates are influenced by the inferred position of fossil samples as sampled ancestors or fossil tips in the phylogeny (Beaulieu and O’Meara, 2023; Barido-Sottani and Morlon, 2025), and we therefore tested the ability of the F-ClaDS model to correctly identify sampled ancestors. Finally, we applied our new F-ClaDS approach to a dataset of Nimravidae, a fully-extinct carnivoraform family that spanned from the Early Eocene to the Late Miocene (Poust et al., 2022). A recent study performed a phylogenetic inference for this group using the homogeneous FBD process (noted HFBD hereafter) as the tree model (Poust et al., 2022). However, we expect that diversification rates are highly heterogeneous in this clade, in particular due to changes in diet. The specialisation of species to a carnivorous diet highly correlates with morphometrical measures on teeth in the mammalian order Carnivora (Valkenburgh, 1993), meaning that we can infer with relative confidence the past diet and the degree of diet specialisation in the fossil record of species in this clade (Van Valkenburgh, 1991). The hypercarnivorous diet (i.e., more than 70% of the diet composed of meat) is especially interesting for diversification dynamics, as the fossil record seems to show iterative radiations of clades developing this ecology, with usually shorter lifespan relative to hypocarnivorous or mesocarnivorous species (Van Valkenburgh, 1991, 2007; Slater, 2015; Balisi et al., 2018). We thus expect that Nimravidae shows a high heterogeneity in rates and that the subclades identified as hypercarnivorous (Hoplophoninae and Barbourofelinae) have higher average diversification rates than the rest of the clade.

## 2 Methods

### 2.1 The ClaDS model with fossils

Similar to the HFBD process, fossil samples are added to the ClaDS model by including a Poisson process with rate *ψ*, capturing the per lineage, per Myr fossilization rate. In the current implementation, *ψ* is set as constant throughout the phylogeny. The F-ClaDS model then uses seven parameters: the speciation and extinction rates at the beginning of the process (*λ*_0_ and *µ*_0_), the parameters characterizing the log-normal distribution in which the new speciation and extinction rates are sampled at each speciation event (*α*_*λ*_, *σ*_*λ*_ for speciation rates and *α*_*µ*_, *σ*_*µ*_ for extinction rates), and the fossilization rate *ψ*. At each speciation event, new birth and death rates are randomly sampled for each daughter branch, so that 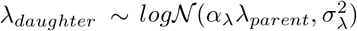 and 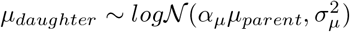.

Our implementation follows the Bayesian data augmentation algorithm described in Maliet and Morlon (2022), building upon its implementation in the ClaDS package in BEAST2 described in Barido-Sottani and Morlon (2023). Bayesian data augmentation samples complete trees (with augmented branches and associated diversification rates, as well as fossilization events in the case of the FBD) during the MCMC and calculates their likelihoods, instead of computing the likelihood of reconstructed trees. New proposals are obtained by proposing new subtrees, as described in Barido-Sottani and Morlon (2023), here simulated under the F-ClaDS process, and accepted according to a Hastings ratio (see below).

To calculate the likelihood of the complete tree under F-ClaDS, the probability that nothing happens along a branch is modified to include the possibility of fossilization events. The probability density of the waiting time *t*_*i*_ between two events on branch **i** then becomes:

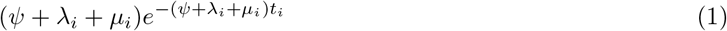

The probability of the event at the end of a branch **i** is 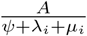 with *A* ∈ {*ψ, λ*_*i*_, *µ*_*i*_} if the event is, respectively, a fossilization, a speciation or an extinction. The probability that no event happened (i.e., on branches leading to present day species) during a time *t*_*j*_ is:

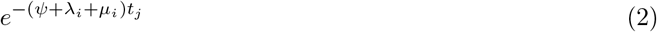

In the complete tree, the number of unsampled extant lineages is known exactly. Noting N the total number of extant lineages and n the number of sampled species in our dataset, the probability of sampling exactly n species is:

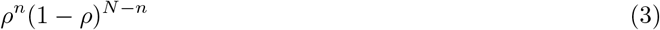

Following Maliet and Morlon (2022), we can note 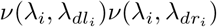 the probability density of the two new speciation rates after a speciation event, 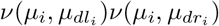 the probability density of the two new extinction rates after a speciation event, *E*_*s*_ the set of internal branches, *E*_*e*_ the set of extinct branches and *E*_*p*_ the set of branches leading to present species. We also note *t*_*i*_ the length of the branch **i**, *E*_*f*_ the set of branches or part of branches (in the case where there is more than one fossil on a branch) leading to a sampled fossil and E the set of all branches. The probability density of the complete tree *T*_*c*_ is:

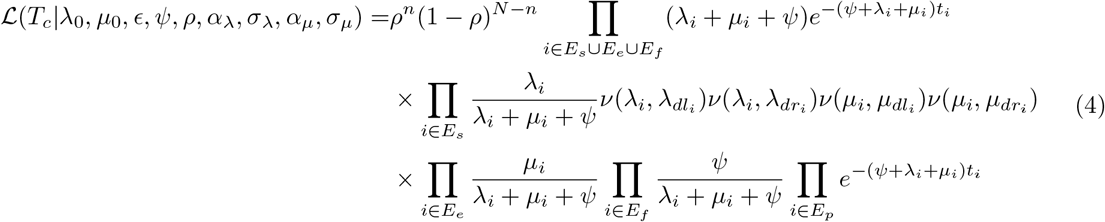

By noting the number of fossil samples *f* and using the log-likelihood as it permits to convert products into sums, which makes the computation easier, we have:

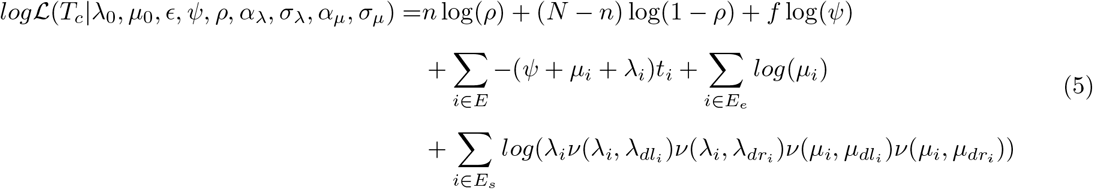

Our implementation in BEAST 2 allows the extinction rate to vary independently of the speciation rate. In what follows however, we fixed the ratio 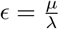 as constant along the tree, in order to reduce computation time (Maliet et al., 2019).

BEAST 2 represents sampled ancestor fossils within trees as tips at the end of 0-length branches, so that all samples are represented as tips (Fig. 1). This creates so-called “fake nodes”, which do not represent speciation events but fossil sampling events, and need to be treated differently in our DA implementation. In particular, *λ*_*i*_ and *µ*_*i*_ are not updated when a sampled ancestor occurs (Fig. 1b), as it is a sampling on a branch and does not correspond to a speciation event. When a fossil has no sampled descendants, it is represented as a true tip. The DA simulates the branch on which it is sampled (Fig. 1a), until this lineage goes extinct (or goes to present but is not sampled).

**Figure 1:**
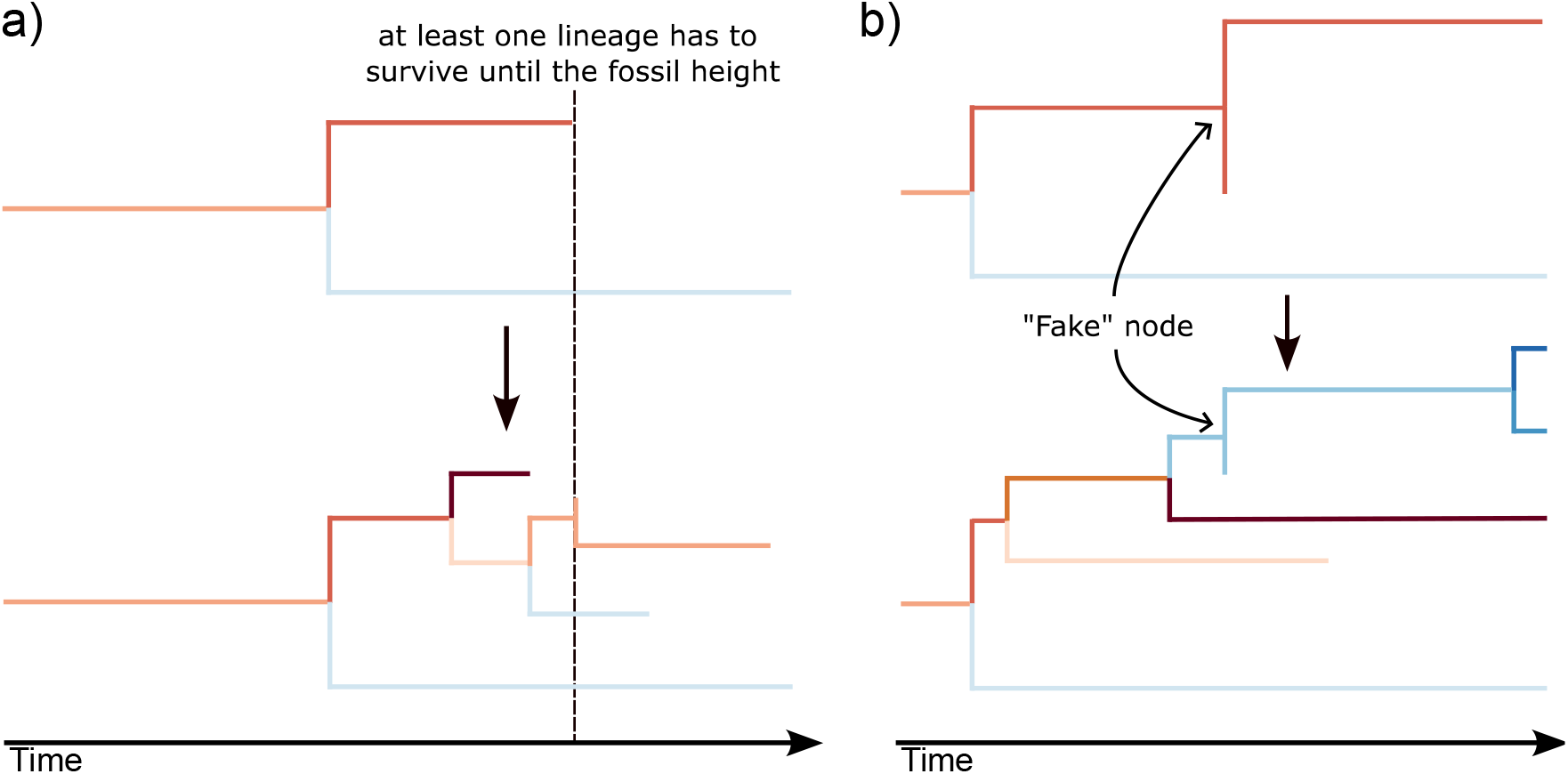
Data Augmentation under different fossil configurations. Top panels show the observed tree, bottom panels show the tree after augmentation. In both cases, the branch on the top, after the split, is the one that is augmented. In both cases, the simulation procedure starts at the node above the fossil sample and continues until reaching the age of the fossil sample. At this time point, we select one lineage at random between all lineages of the augmented tree which exist at the time of the sample. In panel **A**, the fossil taxa is placed as a tip. In this situation, the fossil is placed as a sampled ancestor on the selected lineage, and all lineages then continue the simulation until the present. In panel **B**, the fossil taxa is a sampled ancestor. In this situation, the selected lineage is assigned to match the fossil lineage in the sampled tree, and gets removed from the simulation. All the other lineages continue the simulation until the present. The data augmentation proceeds then to the edge below the fossil sample. In both cases, speciation and extinction rates remain identical after the sampling event. Colors show the variation in the speciation rate, *λ*, along the tree.

During the MCMC search, new augmented trees are proposed following a Metropolis-Hastings algorithm (Höhna and Drummond, 2012). If a new tree *T*_1_ is proposed from an initial tree *T*_0_, the probability of accepting this new state is min 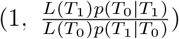 with L(.) the posterior density of the tree (Maliet and Morlon, 2022). After each modification, new augmented trees are simulated along the modified branches, then the ratio 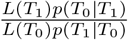 is calculated and the new state is accepted or rejected. If a valid augmented tree cannot be simulated, the proposal is automatically rejected.

As in the SA package that implemented the FBD process, DA operators were modified to be compatible with tips at different heights (because of fossil taxa) and especially to avoid negative-length branches during subtree-swapping, moving or scaling. We also implemented a “Leaf to Sampled Ancestor” operator that changes a fossil tip into a sampled ancestor of its sister lineage and vice-versa (Heath et al., 2014). At each modification of the tree, F-ClaDS updates the augmented tree at the modified branches, so if a branch including sampled ancestors is moved, the rates are also updated for the branches under the fake node created by the fossil. Moreover, we modified the different operators of the inference so that they do not move fossils that are sampled ancestors: once a fossil specimen is set as a sampled ancestor, it is “locked” on this lineage until an operator changes it from a sampled ancestor position to a sister clade position. However, a lineage containing a sampled ancestor can still be moved, the operators are just set so that a sampled ancestor and its descendant node are considered as one lineage only.

Finally, our implementation can include stratigraphic uncertainty for fossils, as previous studies have shown that integrating fossil age uncertainty leads to better prediction of diversification rates (Drummond and Stadler, 2016) and provides more accurate fossil age estimates (Barido-Sottani et al., 2023).

We implemented F-ClaDS as a tree prior in the BEAST2 package ClaDS (Barido-Sottani and Morlon, 2023), and integrated the possibility to run this model through the BEAUti graphical user interface. As such, the implementation is fully compatible with all other functions of the BEAUti interface, such as using multiple alignment partitions, customizing the substitution and clock models, and selecting a starting tree.

### 2.2 Validation

To check that our implementation of F-ClaDS is correct, we used a process of simulation-based calibration (Talts et al., 2018), previously used for other birth-death processes using fossil data (Andréoletti et al., 2022). This method follows six steps, illustrated in Figure 2:

**Figure 2:**
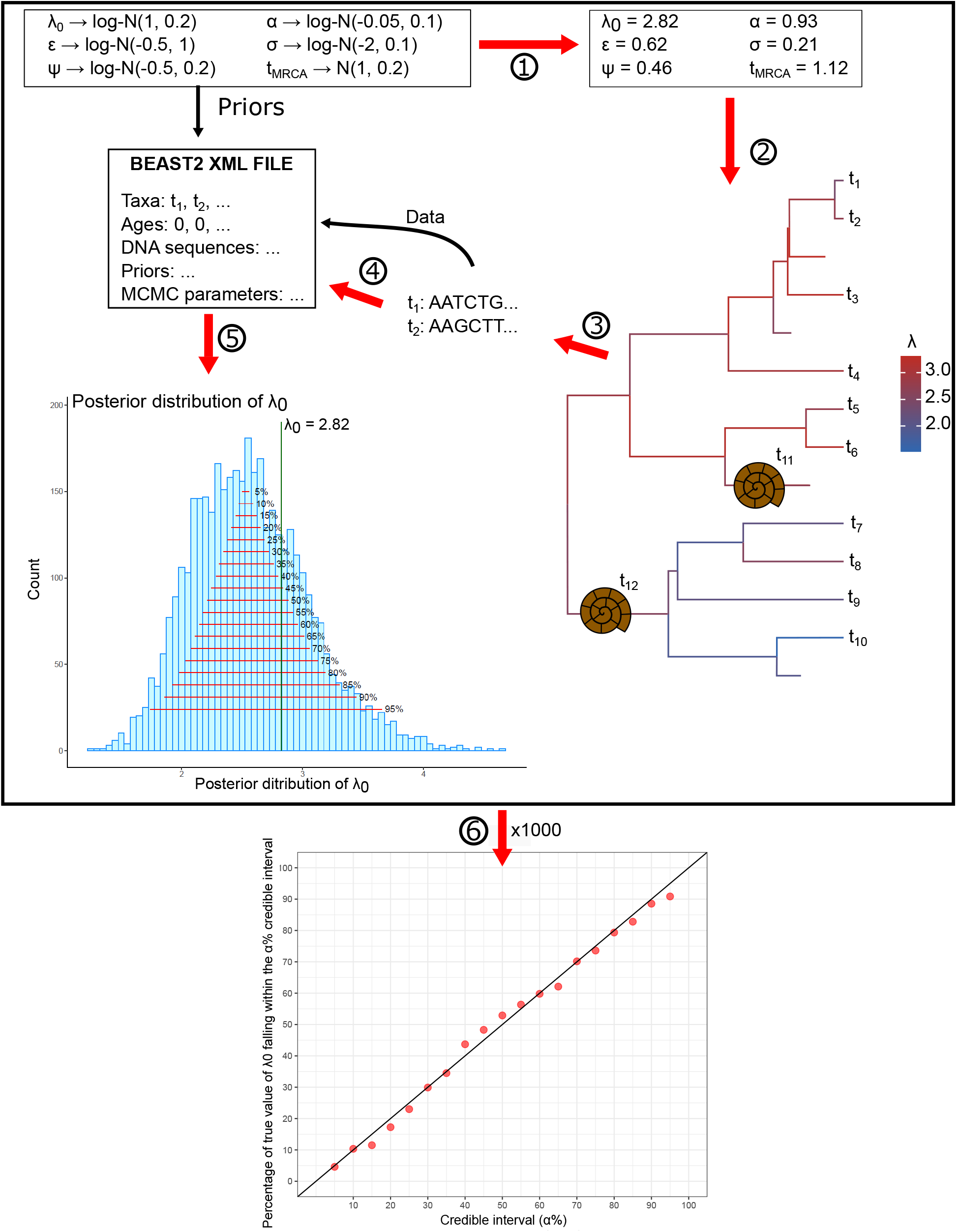
A graphical representation of the validation method. The different steps are detailed in section 2.2: after sampling parameter values from prior distributions (1), we simulate phylogenetic trees including fossils under the F-ClaDS process (2). Next, we simulate sequence alignments on the trees, for both extinct and extant specimens (3). We then use our implementation of F-ClaDS as a tree prior in BEAST2 to infer the tree and posterior distributions of parameter values (4). We check if the true parameter value (green vertical line, illustrated here with *λ*_0_) falls within credible intervals of varying *α*% (5). We repeat the above procedure 1000 times. A good match between the percentage of times when the true value falls in the *α*% credible intervals and *α* validates that the implementation is correct (6).

1. We randomly sample sets of parameters from known distributions (detailed in Table S1). As most empirical studies use a fixed extant species sampling proportion, we fix the value of *ρ* = 0.7.
2. We simulate phylogenetic trees with the generated parameters under F-ClaDS. As the root can be either a branching event or a sampled ancestor, we start the process with a sampled ancestor with a probability 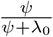.
3. We generate molecular sequences of 500 nucleotides for each sample (fossil or extant) by simulating a Jukes-Cantor process (Jukes et al., 1969) along the simulated phylogeny.
4. We infer the phylogeny and the parameters with BEAST2 using the simulated sequences and the age of the samples as data. The inference uses the same priors and models (including F-ClaDS) as for generating the parameters and sequences.
5. We compare the true values of the parameters used to simulate our data to the posterior distribution obtained from the inference in BEAST2. We compute the credible intervals of *α*% with *α* varying from 5 to 95 by steps of 5, and for each we check whether the true value falls into the *α*% credible interval.
6. We did 1000 simulations following steps 1-5 above, and computed for each *α* the percentage of replicates where the true parameter value was included into the *α*% credible interval.

Our expectation is that if the inference and simulation models match perfectly, then *α*% of the replicates will have the true value fall into the *α*% credible interval, for all values of *α*.

F-ClaDS is also applicable to phylogenies with only fossils. We validated this application as well, following the same validation process as described above, but with the probability of sampling extant lineages to 0 (ie, *ρ* = 0). The prior distributions of all the parameters are detailed in Table S2.

### 2.3 Estimating the number of sampled ancestors

Whether specimens in the fossil record are likely to be sampled ancestors of others (i.e., belonging to the same branch) has been questioned early in the development of cladistic methods (Farris, 1976; Hull, 1979; Paul, 1992). Theoretical work showed that, while very few specimens in the fossil record are described as sampled ancestors of either extant or extinct specimens, the proportion of recovered fossils that are sampled ancestors should be quite high, especially in well-sampled clades (Foote, 1996; Parins-Fukuchi and Saulsbury, 2025). In phylogenetic analyses, the FBD process accounts for this by allowing fossils to switch from a sister clade position to a sampled ancestor position during the inference, therefore integrating sampled ancestors into the phylogeny (Heath et al., 2014). However, it is likely that the model of diversification used for inference can affect the amount of fossil specimens inferred as being sampled ancestors (Barido-Sottani et al., 2020). In particular, if diversification rates are highly heterogeneous but the phylogenetic inference is performed under an assumption of homogeneous rates, fossil specimens belonging to the same branch (and which therefore should be inferred as sampled ancestors) may be misleadingly inferred as fossil tips, to compensate for the unaccounted for rate heterogeneity. We checked this by comparing the ability of both the homogeneous-rates FBD and F-ClaDS to recover the correct number of sampled ancestors when the tree is generated via the F-ClaDS process. We generated phylogenetic trees under the F-ClaDS process with fixed parameters: *λ*_0_ = 1.2, *ψ* = 1, *ϵ* = 0.3 and a root height of 2. We chose those parameters values to generate trees with reasonable numbers of fossils (23 on average, with range from 5 to 124). We then tested different sets of *α* and *σ* values, as those control how much rate variation is present across the tree (*α* ∈ {0.9, 1, 1.1, 1.2} and *σ* ∈ {0.1, 0.2}). We only kept trees with more than 40 tips, and generated 100 trees for each combination of *α* and *σ* values. On each tree, we simulated the evolution of a 500 nucleotide sequence according to a JC69 model. We then used the sequences and ages of each sample to infer the trees, with both the HFBD process and F-ClaDS. Finally, we calculated how many samples were inferred to be sampled ancestors for the two diversification models used in the inference, and compared these numbers to the true values.

### 2.4 The diversification of Nimravidae

We used a dataset from a previous phylogenetic study of Nimravidae, with 36 nimravid species and 4 species as outgroup (Barrett, 2021; Poust et al., 2022) coded in 225 morphological characters. The phylogeny has previously been inferred in BEAST2 with a constant-rate HFBD process as a tree prior. Here, we infer the tree with F-ClaDS as a tree prior. Including outgroups in Bayesian phylogenetic inferences is generally not recommended, as their sampling probability does not match with the probability set for the ingroup. Thus, we removed the 4 outgroups (*Nandinia binotata, Procynodictis vulpiceps, Hesperocyon gregarius* and *Tapocyon robustus*) of the dataset before running the analysis. As Nimravidae does not have any extant representative, we fixed *ρ* = 1. We also ran an analysis with the HFBD by setting *α* = 1 and *σ* = 0 in the ClaDS package in BEAST2. These provided us with augmented trees output for these two analyses, which we used to estimate diversity-through-time curves. In both cases, we used the same prior distributions for *λ*_0_ and *ψ* (Exp(1)), for *ϵ* (U(0, 1)) and for the root age (Exp(1), with a 37 Ma offset). For the F-ClaDS analysis, we also used a prior *log*N (0, 0.05) for *α* and *log*N (−2, 0.1) for *σ*. For the other parameters, we used the same priors as Poust et al. (2022).

For both analyses, we ran 10 independent runs with a MCMC chain length of 100,000,000 operations. We then used a 10% burn-in in the combined files of all the analyses that converged (9 for the F-ClaDS analysis, 8 for the HFBD analysis). To assess the convergence of the whole analysis, we used an Effective Sample Size (ESS) threshold of at least 200 for each parameter of the inference. In this analysis, we recovered high ESS for all the parameters, the minimum ESS being 526 for the F-ClaDS analysis and 358 for the HFBD analysis. Generating the MCC tree is complicated when all the taxa in the analysis are fossils and we are taking into account the stratigraphic uncertainty (Barido-Sottani et al., 2018). We followed the procedure detailed in Barido-Sottani et al. (2018) tutorial for FBD to generate the MCC tree in this case, by adding a “dummy” lineage going from the root to present to all the trees of the posterior, and then estimating the MCC tree on those new trees using the TreeAnnotator tool provided with BEAST2. After that, we remove the dummy tip to get the proper MCC tree. As the age of the tips varies from one tree to another, we used the “Keep target heights” option in the TreeAnnotator software to generate the MCC tree, which avoids having negative branch lengths.

In order to test the hypothesis that the most hypercarnivorous clades *Hoplophoneus* and (*Barbourofelis* + *Albanosmilus*) experienced higher speciation and extinction rates, we computed the distribution of the speciation rates for these groups and compared them to rates on the rest of the phylogeny, using augmented trees. We also used augmented trees to estimate diversity-through-time trajectories for the full phylogeny and the two hypercarnivorous subclades.

## 3 Results

### 3.1 Validation

The results of the validation on trees with extant species are shown in Figure 3, and those on fully extinct phylogenies in Figure S1. The parameters are well estimated during the validation (Fig. 3a and S1a). The tree height is well-estimated when the root corresponds to a speciation event (Fig. 3b), but looks “too well” estimated when it is a sampled ancestor (Supplementary Figure S2). This is likely due to issues with calculating a credible interval while having a hard boundary in the distribution. If the root of the tree is a sampled ancestor, then the true value for the tree height is exactly the age of the oldest fossil. During the inference, the tree height cannot be estimated lower than the age of the oldest fossil, which leads to a one-sided posterior distribution where a large amount of weight is on a single value at the extremity of the range, a situation the credible interval calculation is not designed for. Since the tree height and all the other parameters are well estimated in cases where the first event is a speciation rather than a fossilization, this problem is likely an artifact of the interval construction rather than due to an implementation issue.

**Figure 3:**
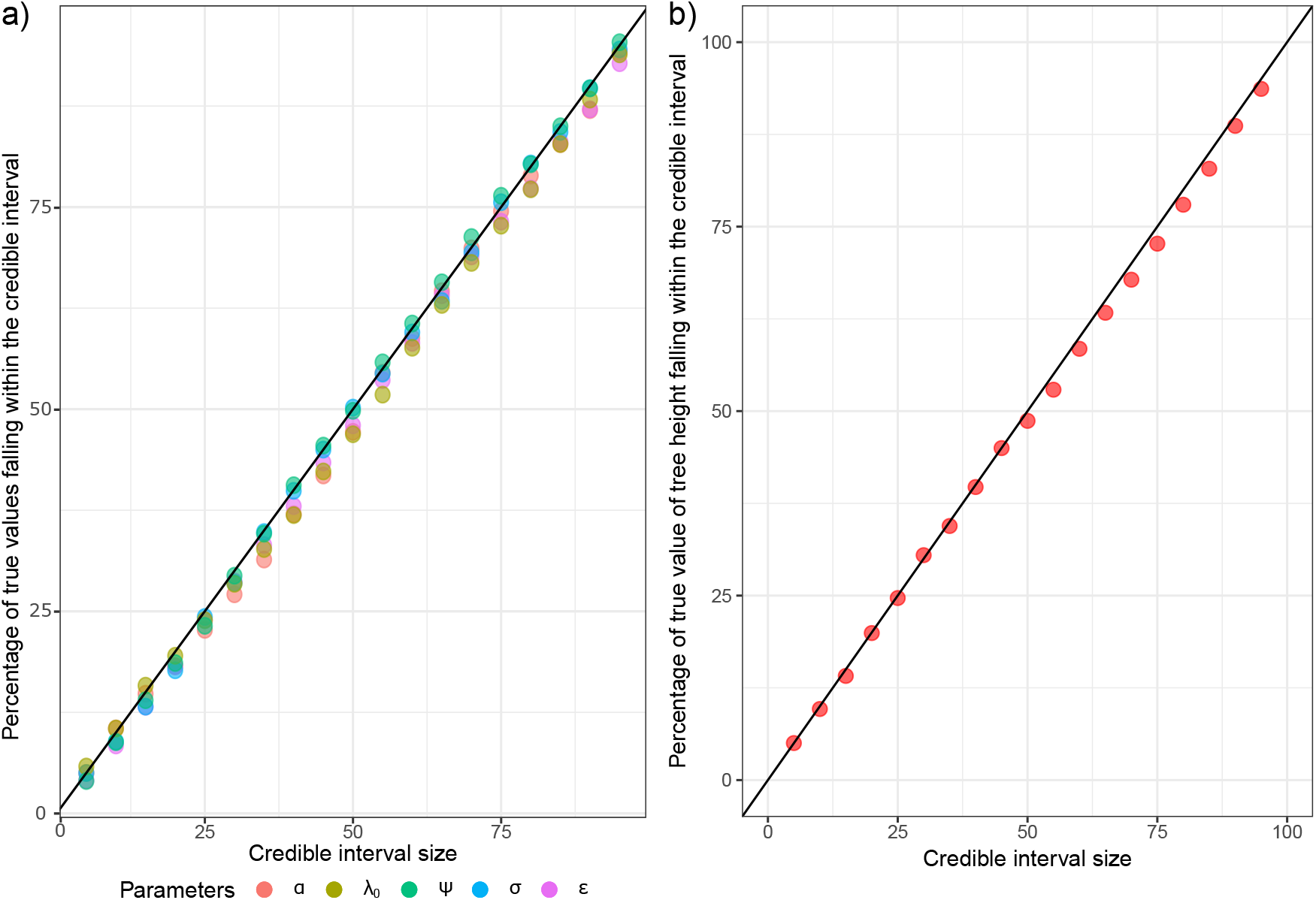
(a) Validation plot for 1000 simulations. The black line represents the 1:1 line and is the aim of the validation. (b) Validation plot for the 758 simulations which started with a speciation event (out of 1000 total, see Section 2.2). The black line represents the 1:1 line and is the aim of the validation.

### 3.2 Estimating the number of sampled ancestors

The generated trees had on average 66.9 tips and 23.7 fossils (either tips or sampled ancestors), 21.7 of which were in a sampled ancestor position. We show the absolute difference between the estimated and true number of sampled ancestors in Figure 4 for *σ* = 0.1, as the results are similar to the ones with *σ* = 0.2 (Supplementary Figure S3). We can see that the homogeneous FBD tends to under-estimate the number of sampled ancestors more and more as the values of *α* are moving away from 1 (i.e., as the rates are becoming more and more heterogeneous), with, on median, 1.1 less sampled ancestors identified as such with the constant-rate FBD compared to the true value. Inferences under the F-ClaDS model are mostly unbiased, with a median estimate very close to the true value for all values of *α*.

**Figure 4:**
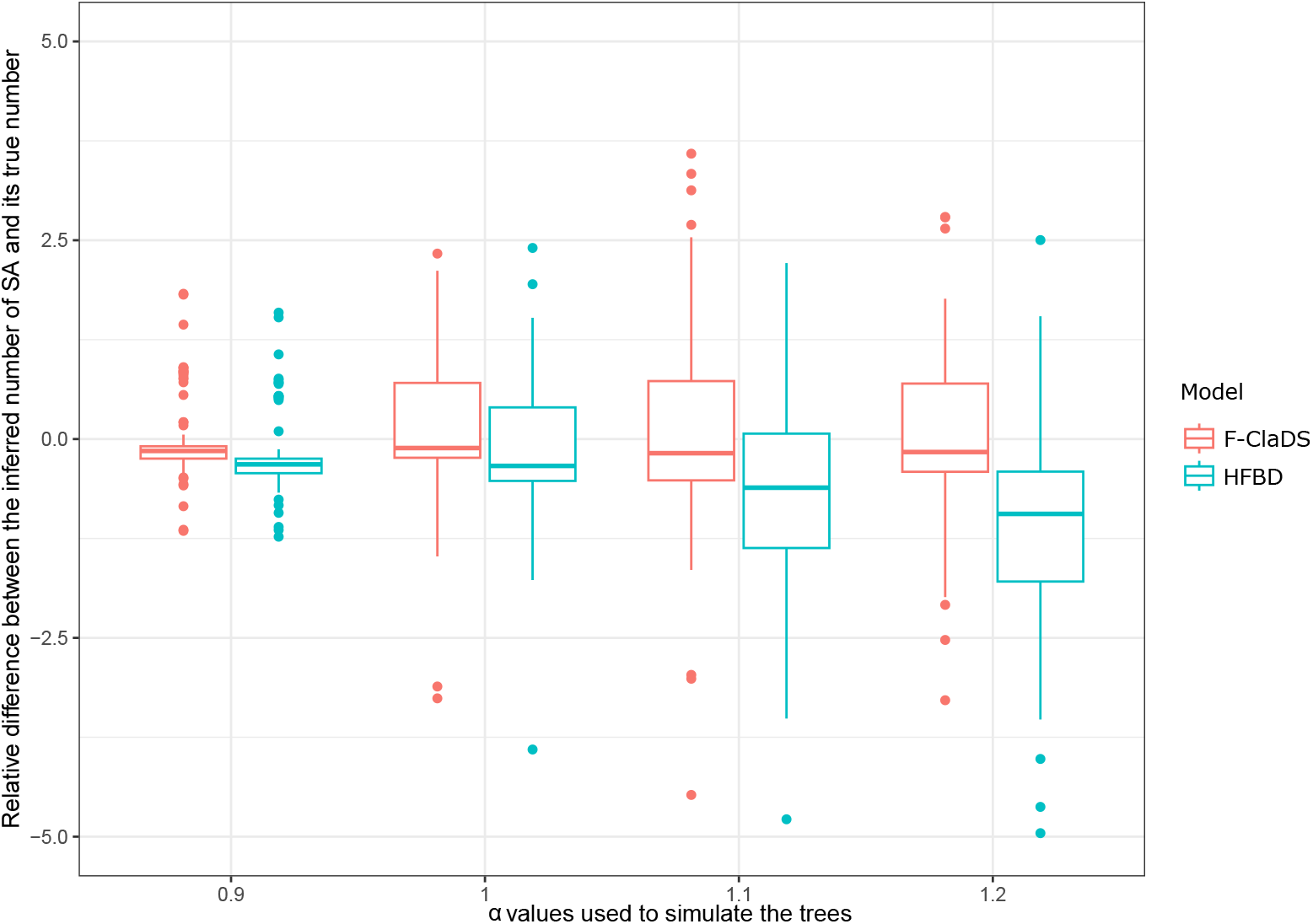
Boxplot of the differences between the number of sampled ancestors estimated during the inference under both F-ClaDS and the homogeneous FBD and the true number of sampled ancestors, for different values of *α* with *σ* = 0.1. Outliers are not shown, as extreme values were collapsing the boxes.

While both *α* and *σ* affect the heterogeneity of the rates across the tree, only *α* seems to influence the mismatch in estimated number of sampled ancestors, while increasing *σ* did not affect the difference in the inferred number of sampled ancestors. This may be explained by the fact that *σ* affects the variance of the rates across the tree, but does not affect the expected values of the rates. On the other hand, increasing *α* away from 1.0 leads to descendant rates which are on average more different from the ancestor rate, and thus to higher overall heterogeneity in the process.

### 3.3 The diversification of Nimravidae

The topologies of the two MCC trees obtained with either the F-ClaDS or the HFBD model as a tree prior were very similar, but the branch lengths and the ages of nodes and tips varied quite a lot (Fig. 5a,b). The basic relationships between the taxa did not change substantially. In particular, the *Nimravus* genus was recovered as paraphyletic in both analyses. There were two differences in parts of the tree with low posterior node support (Fig. 5a,b): i) *Nimravus brachyops* was recovered as the sister taxa of *Dinaelurus crassus* in the F-ClaDS analysis, with a low posterior support (0.02), while *D. crassus* was recovered as an independant lineage in the HFBD analysis; ii) *Albanosmilus jourdani* was recovered as the sister taxa of (*Barbourofelis piveteaui* +*Barbourofelis morrisi*) in the HFBD analysis, while it was within a clade including *A. whitfordi, B. loveorum* and *B. fricki* in the F-ClaDS analysis. In terms of node and tip ages, the root age was recovered as older with the ClaDS (39.56 Ma, 95% HPD interval [37.07, 42.83]) versus HFBD (38.52 Ma, 95% HPD interval [37.01, 40.62]) analyses. Also, the F-ClaDS analysis recovered on average a higher number of sampled ancestors in the observed tree, with a mean of 14.3 versus 13.7 for the HFBD analysis (Fig 5c).

**Figure 5:**
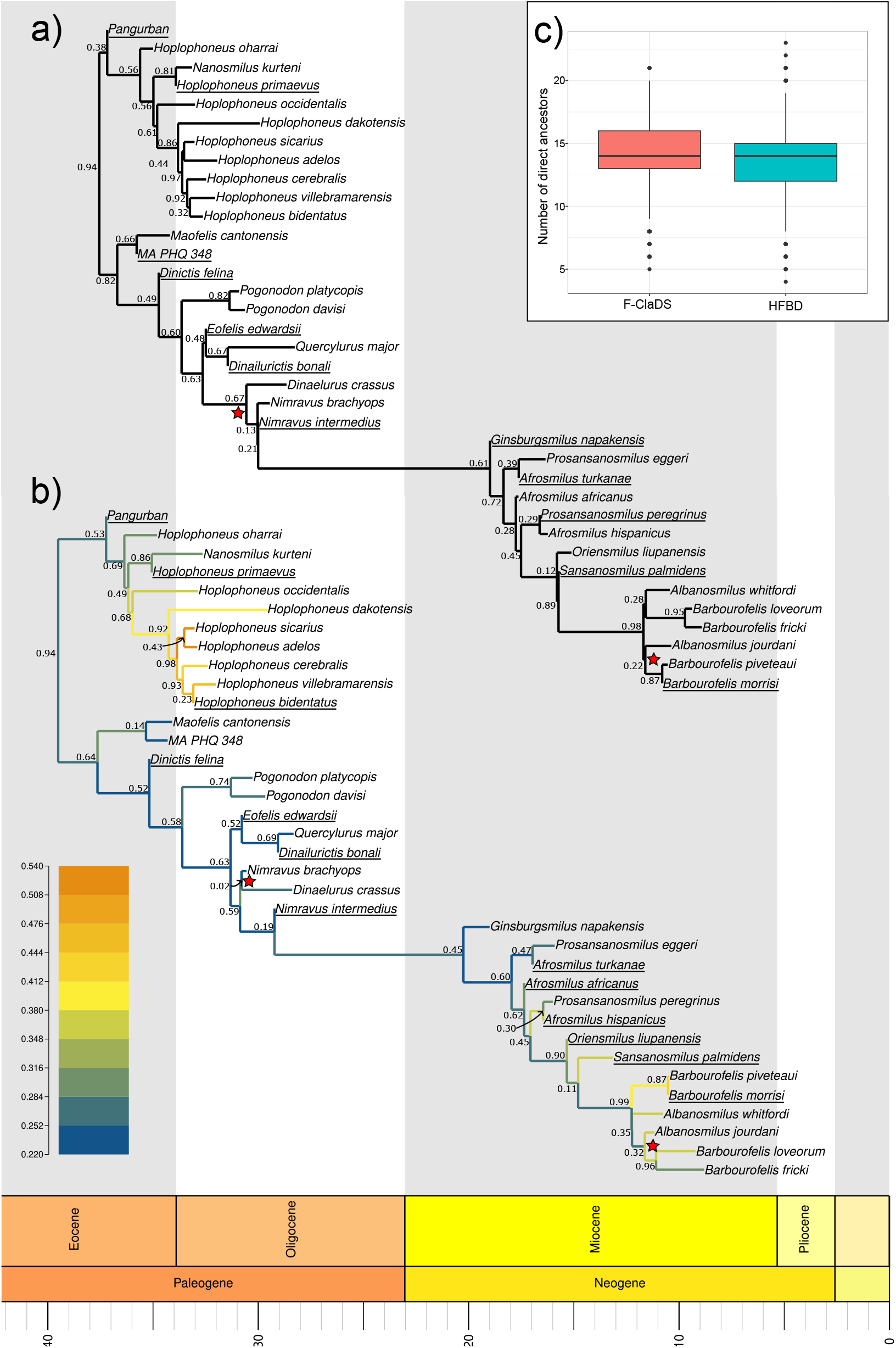
MCC trees of Nimravidae obtained with (a) a homogeneous FBD as a tree prior and (b) with F-ClaDS as a tree prior. Branch colors indicate the median speciation rate in the posterior. Values at the internal nodes show their posterior support, red stars indicate the parts of the topology that are different in both analyses. c) Distribution of the number of sampled ancestors averaged over respectively 8109 and 7208 trees from the posterior distribution of the Nimravidae’s observed tree, with the two tree prior models.

We recovered an average per lineage speciation rate of 0.47 Ma^*−*1^ (95% HPD interval [0.215,0.771]) in the HFBD analysis, while we recovered increasing speciation rates in the F-ClaDS analysis, with *λ*_0_ = 0.26 Ma^*−*1^ (95% HPD interval [0.076,0.506]) and *α* = 1.048 (95% HPD interval [0.953,1.144]), and strong contrasts between lineages (*λ* ∈ {0.22, 0.54} in the MCC tree). We recovered on average *σ* = 0.129 (95% HPD interval [0.104,0.157]), giving an average expected relative change in speciation rate 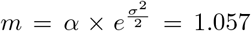. The fossilization rates were in the same range in both analyses, with an average value *ψ* = 0.258 for the ClaDS analysis (95% HPD interval [0.126,0.402]) and *ψ* = 0.280 (95% HPD interval [0.137,0.435]) for the HFBD analysis. In particular, the diversification rates were recovered as high within the *Hoplophoneus* genus (median value of *λ* = 0.432, 95% HPD interval [0.177,1.757]) and within the (*Barbourofelis* + *Albanosmilus*) complex (median value of *λ* = 0.380, 95% HPD interval [0.140,1.962]), while being low for the rest of the clade (median value of *λ* = 0.264, 95% HPD interval [0.129,1.617], see Supplementary Figure S4). We also recovered a slightly higher turnover on average for the F-ClaDS analysis, although the difference is not significant, with a mean estimate *ϵ* = 0.923 (95% HPD interval [0.7954,1]) versus *ϵ* = 0.883 (95% HPD interval [0.7133,1]) for the HFBD analysis.

The inferred diversity-through-time plot shows a peak of diversity just after the Eocene/Oligocene transition, with an estimated average diversity of 13 species, mainly due to the diversity of the *Hoplophoneus* genus (Fig 6b), followed by a decrease in diversity throughout the Oligocene with an estimated diversity of 2.5 species around the Oligocene/Miocene transition (Fig. 6a). Afterward, there is a small increase in the number of lineages around 19-15 Ma during the Miocene Climate Optimum (Steinthorsdottir et al., 2021). Another peak of diversity is recovered around 10 Ma, with an inferred average of 5 species, followed by a complete loss of diversity during the Late Miocene and Pliocene. These patterns are recovered in both analyses, but diversity varies slightly less in the HFBD analysis (Supplementary Figure S5).

**Figure 6:**
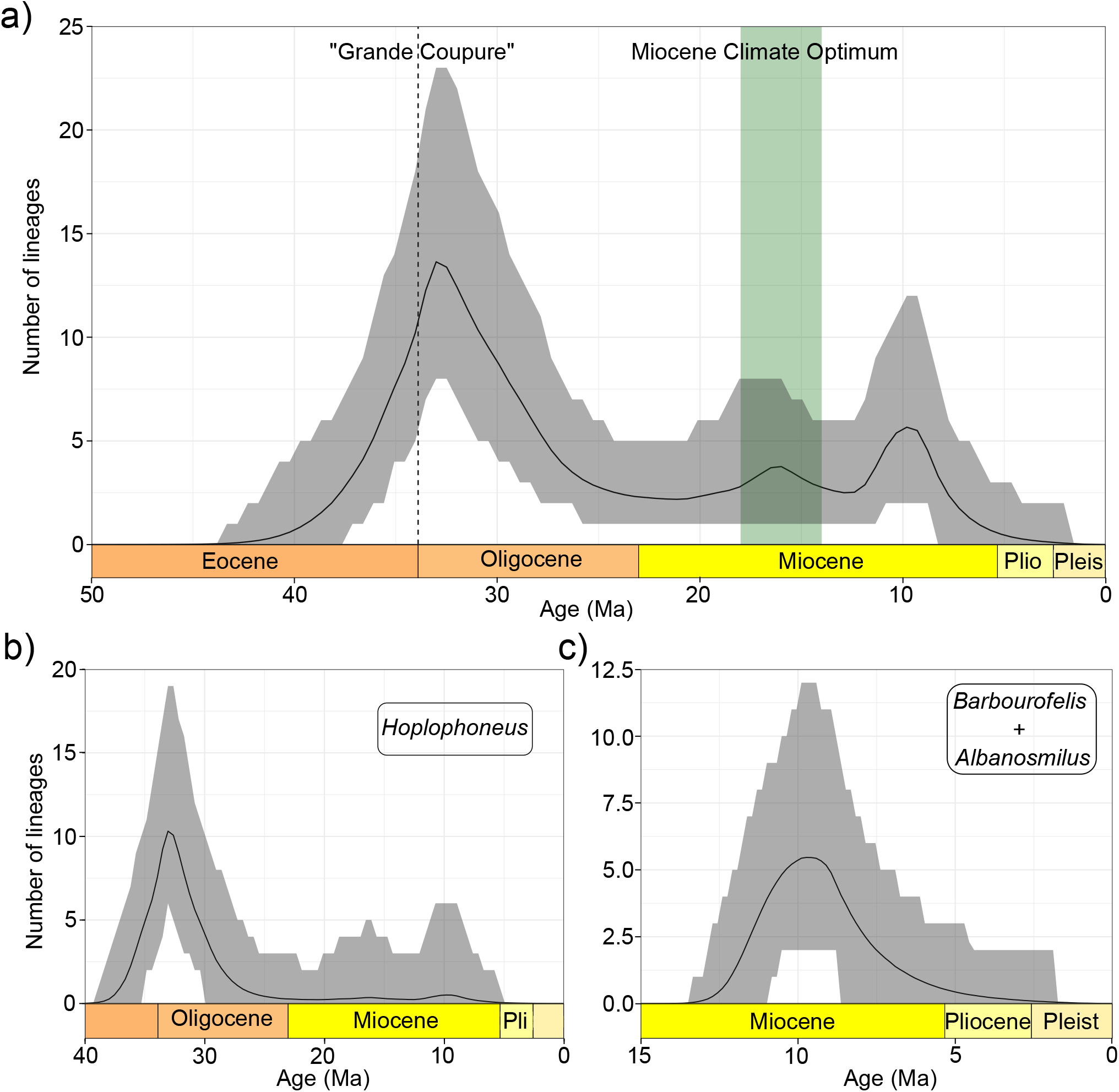
Estimated diversity-through time, based on the posterior distribution of the augmented trees, when the tree prior model is F-ClaDS for (a) all nimravids, (b) the *Hoplophoneus* genus and (c) the (*Barbourofelis* + *Albanosmilus*) complex. The full black line shows the mean value, the grey area shows the 95% credible interval.

## 4 Discussion

We have built and validated a model combining the FBD process, which allows phylogenies to integrate samples from the fossil record, and the ClaDS process, which models progressive changes in diversification rates across a phylogeny; we have integrated this model in the ClaDS BEAST2 package. Our combined F-ClaDS model provides a nuanced model for the diversification process of phylogenies combining extant and extinct samples. The model can be run on a fixed tree, but since it is fully integrated into the BEAST2 framework, it can also be easily combined with other components, such as a molecular evolution model and the Mk Lewis morphological evolution model, as we demonstrate in our empirical application on nimravids.

We showed that heterogeneity in the diversification rates can lead to under-estimate the number of sampled ancestors during the phylogenetic inference when using the constant-rates homogeneous FBD process as a diversification model. We demonstrated this by simulation, and the tendency was confirmed in our empirical analysis, where using F-ClaDS as a tree prior lead to a slightly higher estimated number of sampled ancestors in the Nimravidae tree than when using the HFBD (Fig. 5c). The effect of heterogeneous rates on the inference of the number of sampled ancestor is particularly interesting as many ecological features may lead to variation in the diversification rates across the tree, for example in the case of adaptive radiations (Stroud and Losos, 2016), or when speciation or extinction are influenced by phenotypical traits that vary across lineages (Purvis, 2008). The widespread use of constant-rate homogeneous FBD as a tree prior could thus contribute to the apparent underestimation of the actual number of sampled ancestors in phylogenetic analyses (Parins-Fukuchi and Saulsbury, 2025). Another potential (larger) contributor to this discrepancy is the mismatch between the assumption of FBD models, including F-ClaDS, that fossil samples represent individual occurrences sampled according to a Poisson process, and empirical datasets, including our empirical Nimravidae dataset, composed of morphospecies with often a single representative fossil specimen used in the phylogenetic inference. Modeling morphospecies ranges in F-ClaDS, for instance by integrating the FBD-range model (Stadler et al., 2018), or accounting for multiple occurrences of fossils matched to morphospecies in paleodatabases (Mitchell et al., 2019), will be needed to fully correct biases in the inference of sampled ancestors and associated biases in the inference of fossilization and diversification rates.

In our empirical dataset, accounting for rate variations by using the F-ClaDS model as a tree prior did not have a major impact on the topology of the tree, but it strongly impacted estimated node and tip ages, and thus branch lengths. The slight differences observed in the topology, for example in the *Albanosmilus* and *Barbourofelis* genera, affected species whose placement in the topology is quite uncertain (Fig. 5a,b). For example, *A. whitfordi* has only recently been moved from the *Barbourofelis* genus to the *Albanosmilus* one (Robles et al., 2013). Unlike the previous analysis of this dataset, F-ClaDS also allows us to identify variations in the diversification process between the different lineages. In particular the *Hoplophoneus* genera and the (*Barbourofelis* + *Albanosmilus*) complex are estimated to have elevated speciation and extinction rates (Supplementary Figure S**??**). While the use of a constant turnover through time in our analysis enforces speciation and extinction rates to rise or fall together, previous studies also found high speciation and extinction rates in the most specialised nimravids (Silvestro et al., 2015; Balisi and Van Valkenburgh, 2020), showing that our results are consistent with previous studies without such constraint on the turnover.

The quick extinction of the two hypercarnivorous clades *Hoplophoneus* and (*Barbourofelis* + *Albanosmilus*) match previous observations in other hypercarnivorous group such as canids, and has been explained either by their sensitivity to environmental change, due to their high position in the trophic chain, or by active out-competition by other carnivorous clades (Van Valkenburgh et al., 2004; Van Valkenburgh, 2007; Balisi et al., 2018; Balisi and Van Valkenburgh, 2020). The latter hypothesis is the most supported in the case of canids (Silvestro et al., 2015), in which when an hypercarnivorous group goes extinct, another less carnivorous clade replaces it in the hypercarnivorous niche, with Eocene/Oligocene Hesperocyoninae being replaced by Borophaginae during the Miocene, themselves replaced by Caninae during the Pliocene and Pleistocene. For nimravids, the first known hypercarnivorous species is *Pangurban eigiae* (Poust et al., 2022), which lived during the Middle Eocene climatic optimum. We estimate a sharp rise in nimravid diversity after this climatic optimum, exemplified by the diverse *Hoplophoneus* genus in the Late Eocene/Early Oligocene (Fig. ‘6b). This period is also marked by huge ecosystem restructuring due to the rapid cooling of the Earth (Liu et al., 2009) at the E/O boundary. It is interesting to note that this major climatic event did not affect nimravids that much, but likely precipitated the extinction of other hypercarnivorous carnivoramorphs such as machaeroidine Creodonta (Zack et al., 2022), and the nimravid hoplophonines filled the hypercarnivorous niche left empty after this extinction event. Following this climate change, we observe a long decline in diversity for nimravids throughout the Oligocene (Fig. 6a,b). We estimate a small diversification event during the Miocene Climate Optimum, a period during which temperature increased, with a peak of mammalian diversity especially in tectonic active regions (Finarelli and Badgley, 2010; Smiley et al., 2024). There is also a second major diversification of hypercarnivorous nimravids, the African barbourofelins, in the Middle/Late Miocene (Fig. 6c), triggered by a complete turnover of the African nimravids, likely due to immigrant taxa from Eurasia (Werdelin, 2022). Silvestro et al. (2015) found that the diversification of barbourofelins and other clades such as Caninae and Felinae likely contributed to the extinction of the Borophaginae. Barbourofelins then went extinct, likely due to the Late Miocene cooling and aridification that, like the Eocene-Oligocene transition, implied large restructurings in the ecosystem (Herbert et al., 2016; Werdelin, 2022). The hypercarnivorous niche was then filled by Caninae and Felidae.

In summary, our empirical application demonstrates that F-ClaDS can leverage the fossil record to accurately recover variations in the diversification process, and confirms that these variations leave evidence in the phylogeny. More generally, allowing diversification rates to vary when using fossil data provides us with a powerful tool to study the radiation of extinct groups with a large disparity in ecology. For example, the diversification of dinosaurs has been largely studied (Benton et al., 2014; Bernardi et al., 2018; Bonsor et al., 2020), especially regarding its dynamic prior to the KPg boundary. The use of F-ClaDS in this case could bring new relevant insights into clade-specific dynamics, as this group experienced a large variety of ecological niches and had a world-wide distribution (Chiarenza et al., 2019). Other groups could be of particular interest, for example the early mammals, where multiple radiations (Grossnickle et al., 2019) are correlated with ecological diversification. Overall, F-ClaDS shows tremendous potential in identifying fine-grained variations in diversification processes throughout the Tree of Life, particularly in the deep past or for completely extinct groups, where most or all of the information is contained in the fossil record. However, phylogeny-based diversification analyses require high-quality input data: well-defined time-calibrated phylogenies are needed to work on a fixed tree, or alternatively large molecular and/or morphological alignments can be used to co-estimate the time-calibrated phylogeny and the diversification rates in a tripartite analysis (Wright et al., 2022). This raises problems for numerous fossil groups, for example sharks (Guinot et al., 2012), for which almost only teeth are available. In this case, ecological and biogeographical features of the clade can be inferred, but not accurate phylogenetic relationships between individual fossil species. Bayesian inference is designed to account for this uncertainty and will reflect it in the distribution of the resulting estimates, but this is likely to result in very large credible intervals from which definitive conclusions will be hard to draw. In addition, F-ClaDS (and most FBD processes, but see Barido-Sottani and Morlon (2025)) currently make the assumption that the fossil sampling rate is homogeneous across the phylogeny. This can induce bias, as some regions of the world are under-sampled compared to others (for example, the fossil record is much better known in North America and Europe than in Africa) and as the possibility of fossilization varies across time in certain regions, as observed in North-American dinosaurs (Chiarenza et al., 2019). Extending F-ClaDS to relax the assumption of a constant fossilization rate will be necessary to more accurately represent the fossilization dynamics through time and across lineages.

Finally, one strong limitation of our current implementation is the computation time required for convergence. Bayesian inference is known to be expensive, and complex models such as F-ClaDS only compound this issue. Our data augmentation approach improves the performance of the model considerably, however the current version of ClaDS remains limited to datasets with around 100 species, which is fairly small especially when including fossil species. Further developments of the model will thus focus on improving its performance to make it possible to apply to larger-scale datasets.

## 5 Conclusion

We extended ClaDS, a diversification model with heterogeneous rates, to combined phylogenies including both extant and fossil data. Our new F-ClaDS model is implemented in BEAST2, a Bayesian inference software which allows to simultaneously infer the time-calibrated phylogeny and the diversification dynamics of a clade. The development, based on data augmentation, involved modifications to the augmented simulations, the likelihood function and the operators, including the addition of a new operator specific to the FBD process. We validated our approach using simulations, and showed that the use of a homogeneous FBD process as the tree prior when diversification rates are heterogeneous leads to an under-estimation of the number of sampled ancestors during the phylogenetic inference. We applied F-ClaDS to a dataset of fossil carnivoraforms, the nimravids, and showed that this tree prior does not have a major impact on the tree topology when compared to the homogeneous FBD model, but does have an impact on fossil and tip ages, as well as on the recovery of sampled ancestors. We recovered higher rates of speciation and extinction in the most specialised hypercarnivorous clades, the hoplophonines and the barbourofelins. This result is in line with previous observation of higher speciation and extinction rates in hypercarnivorous clades, such as extinct canid families. This shows that F-ClaDS is a good tool to test macroevolutionary hypotheses on past diversification processes without a prior knowledge on those processes, and could potentially be used on many empirical datasets, especially fully extinct clades. Several improvements of the model are currently being worked on, such as relaxing the hypothesis of an homogeneous fossilization rate across the tree or reducing the computation time of the model, which currently sets limits to its use.

## 6 Acknowledgements

JBS was supported by the European Union’s Horizon 2020 Research and Innovation Programme under the Marie Sklodowska-Curie grant agreement No. 101022928.

## 7 Data availability

F-ClaDS was added to the ClaDS package starting from version 1.2.0 (for BEAST 2.6) and 2.1.0 (for BEAST 2.7). The package is available for BEAST 2.6 and BEAST 2.7 through the package manager (see installation instructions here: https://www.beast2.org/managing-packages/) and as a public Git repository (https://bitbucket.org/bjoelle/clads/). The R code used for simulation and validation, the BEAST2 XML files used to run the validation and empirical example, and the results of the validation are available on Data Dryad: http://datadryad.org/share/0YPGzUFvQoJumnoXS4Jf4aDpiqBBe2eZbnBT9a1deF4.

