## Supplementary Materials for "The Fossilized Birth Death Process with heterogeneous diversification rates unravels the link between diversification and specialisation to a carnivorous diet in Nimravidae (Carnivoraformes)"

#### 1 Validation

The distribution of the sampled parameters for the validation (Section 2.2) are indicated in Table S1. We fixed the sampling proportion of extant species as $\rho = 0.7$ for all simulations. The distribution of the sampled parameters for the validation with only fossils species is different, to generate trees with enough species. They are indicated in Table S2. To only have fossil species, we fixed the sampling proportion of extant species as $\rho = 0$ for all simulations. Supplementary Figure S1 shows that the validation properly works in this context also.

Supplementary Figure S2 shows the validation plot for the tree height, it shows how the hard boundary set by the oldest fossil can mislead the method used for simulation, as the credible interval built are biased by it.

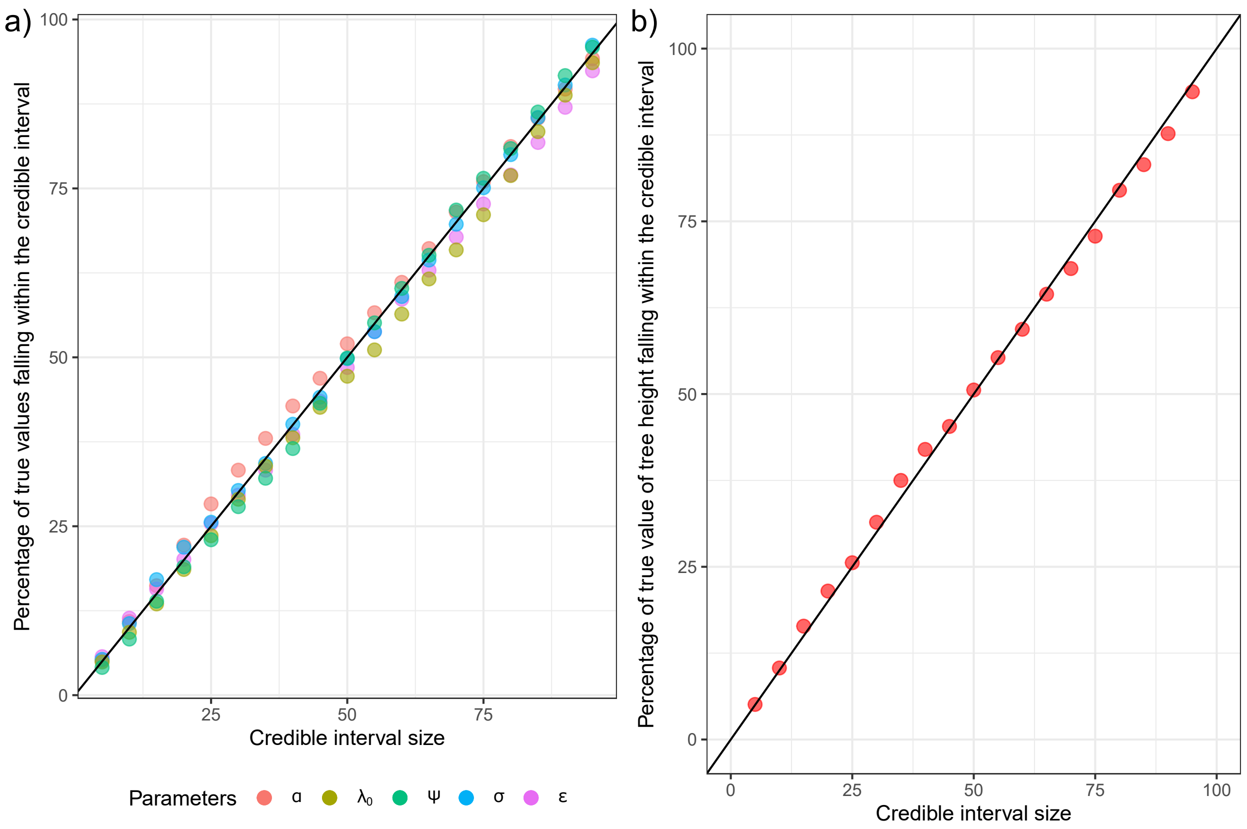


Supplementary Figure S1: Validation plots of simulations with only fossils ($\rho = 0$) for (a) 1000 simulations for all parameters except the tree height. The black line represents the 1:1 line and is the aim of the validation. (b) Validation plot for the 512 simulations which started with a speciation event (out of 1000 total). The black line represents the 1:1 line and is the aim of the validation.

| Parameter | $\lambda_0$ | $\alpha$ | $\sigma$ | $\psi$ | $\epsilon$ | Tree height |
|---|---|---|---|---|---|---|
| Distribution | $log\mathcal{N}(1, 0.2)$ | $log\mathcal{N}(-0.05, 0.1)$ | $log\mathcal{N}(-2, 0.1)$ | $log\mathcal{N}(-0.5, 0.2)$ | $log\mathcal{N}(-0.5, 1)$ | $\mathcal{N}(1, 0.2)$ |

Table 1: Parameter distributions used in the simulations of trees for the validation with extant species ($\rho = 0.7$).

| Parameter | $\lambda_0$ | $\alpha$ | $\sigma$ | $\psi$ | $\epsilon$ | Tree height |
|---|---|---|---|---|---|---|
| Distribution | $log\mathcal{N}(1, 0.2)$ | $log\mathcal{N}(-0.05, 0.1)$ | $log\mathcal{N}(-2, 0.1)$ | $log\mathcal{N}(1, 0.2)$ | $log\mathcal{N}(-1, 1)$ | $\mathcal{N}(1, 0.2)$ |

Table 2: Parameter distributions used in the simulations of trees for the validation with only fossils ($\rho = 0$).

### 2 Estimating the number of sampled ancestors

The Supplementary Figure S3 shows how the estimated number of sampled ancestor varies compared to its true value when we use F-ClaDS as a tree prior or the homogeneous Fossilized Birth Death model. In those simulation, we set $\sigma = 0.2$ when simulating the trees.

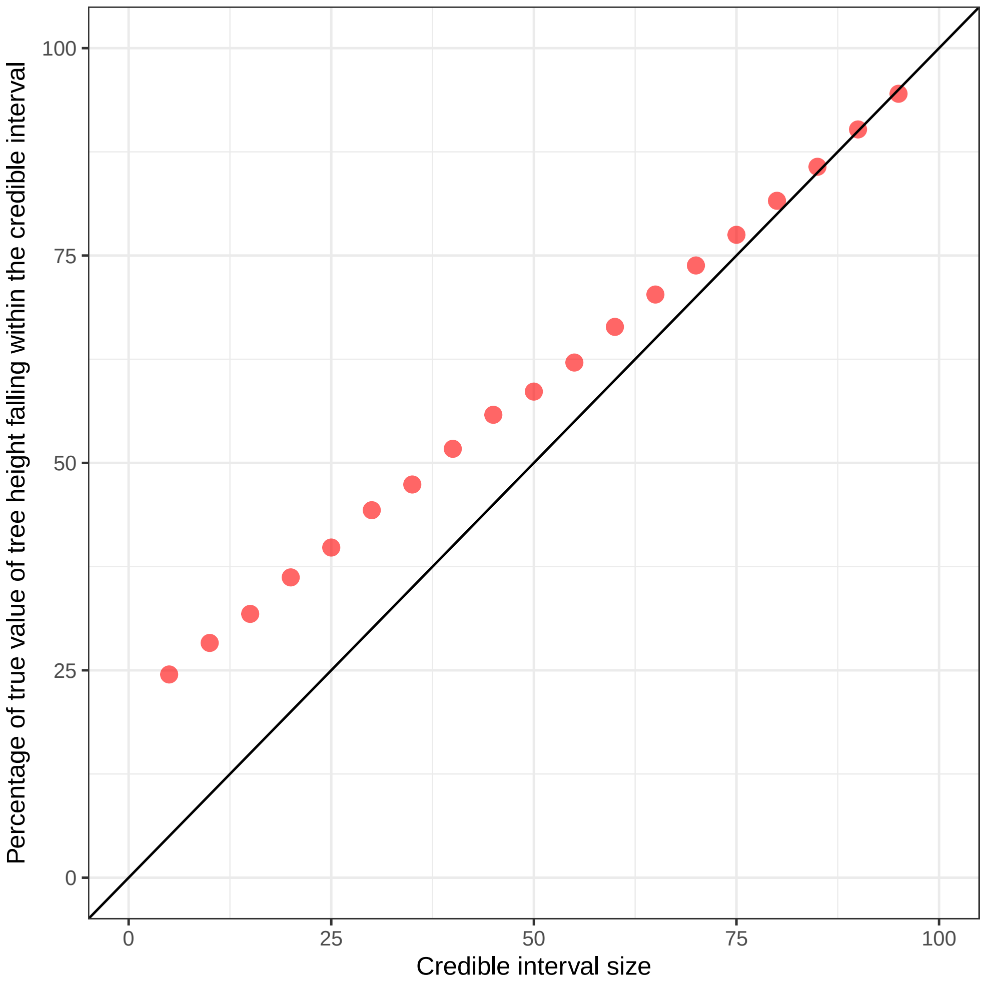


Supplementary Figure S2: Validation plot for 1000 simulations for the tree height. The black line represents the 1:1 line and is the aim of the validation.

### 3 The diversification of Nimravidae

The mean estimates of speciation rates at the tip branches for the (*Barbourofelis*+*Albanosmilus*) complex, the *Hoplophoneus* genus and the rest of the clade were done across 8109 augmented trees from the posterior distribution, and are shown in Supplementary Figure S4.

The lineage-through-time plot for Nimravidae when the HFBD model is used as a tree prior is shown in Supplementary Figure S5. It shows a similar trend as the inference when F-ClaDS is the tree prior (Fig. 6a in the main text), with a peak of diversity right after the Grande Coupure, a smalll increase at the Miocene Climate Optimum and another one with the diversification of barbourofelines around 10 Ma, followed by the extinction of the clade at the end of the Miocene.

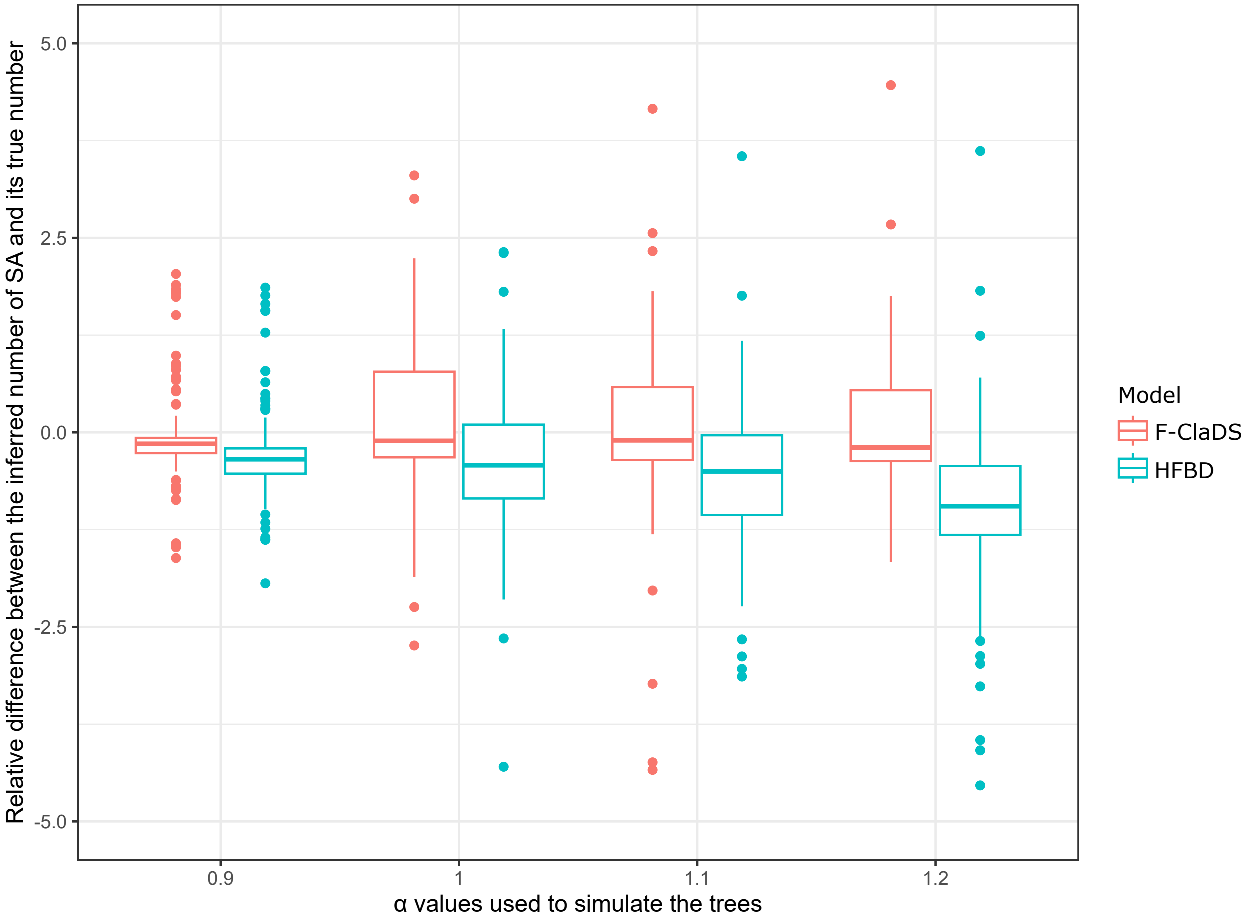


Supplementary Figure S3: Boxplot of the differences between the number of sampled ancestors estimated during the inference under both F-ClaDS and the homogeneous FBD and the true number of direct ancestors, for different values of $\alpha$ with $\sigma = 0.2$. Outliers are not shown, as extreme values were collapsing the boxes.

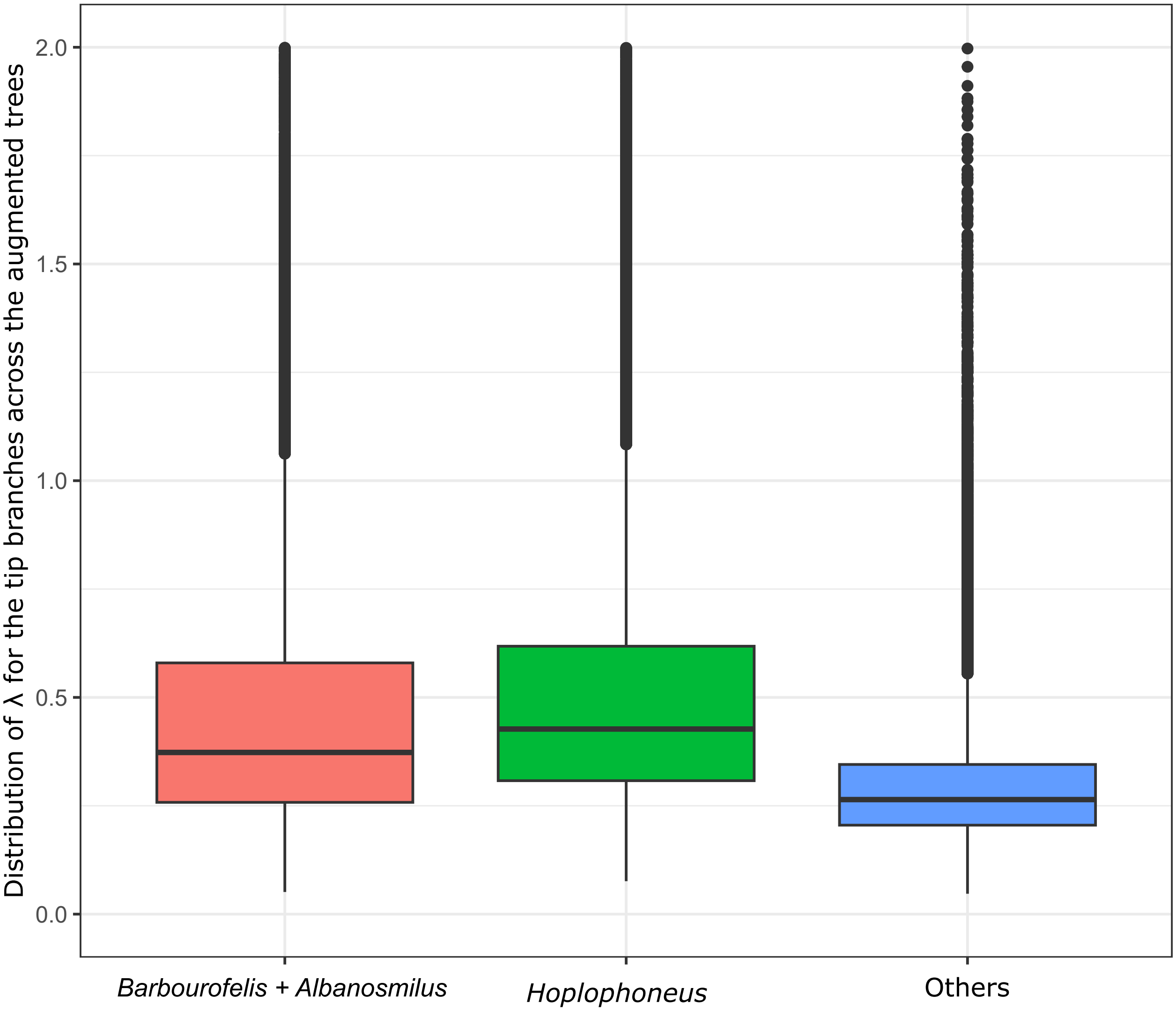


Supplementary Figure S4: Distribution of $\lambda$ values of the tip branches of the two main hypercarnivorous groups and of the other nimravids across 8109 augmented trees from the posterior distribution.

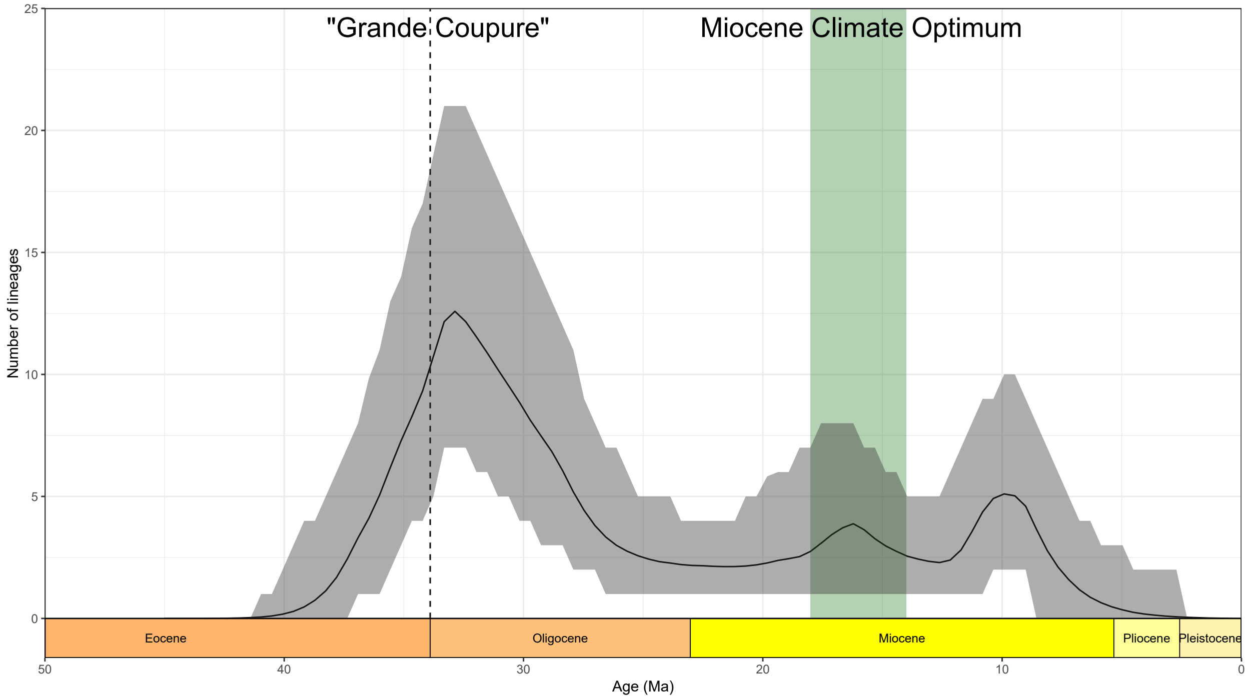


Supplementary Figure S5: Estimated diversity-through time, based on the posterior distribution of 8109 augmented trees, when the tree prior model is the homogeneous birth death process. The full black line shows the mean value, the grey area shows 95% credible interval.
